# Spectroscopic and QM/MM studies of the Cu(I) binding site of the plant ethylene receptor ETR1

**DOI:** 10.1101/2022.06.13.495978

**Authors:** George Cutsail, Stephan Schott-Verdugo, Lena Müller, Serena DeBeer, Georg Groth, Holger Gohlke

## Abstract

Herein, we present the first spectroscopic characterization of the Cu(I) active site of the plant ethylene receptor ETR1. The X-ray absorption (XAS) and extended X-ray absorption fine structure (EXAFS) spectroscopy presented here establish that ETR1 has a low-coordinate Cu(I) site. The EXAFS resolves a mixed first coordination sphere of N/O and S scatterers at distances consistent with potential histidine and cysteine residues. This finding agrees with the coordination of residues C65 and H69 to the Cu(I) site, which are critical for ethylene activity and well-conserved. Further, the Cu K-edge XAS and EXAFS of ETR1 exhibit spectroscopic changes upon addition of ethylene that are attributed to modifications in the Cu(I) coordination environment, suggestive of ethylene binding. Results from umbrella sampling simulations of the proposed ethylene binding helix of ETR1 at a mixed quantum mechanics/molecular mechanics (QM/MM) level agree with the EXAFS fit distance changes upon ethylene binding, particularly in the increase of the distance between H69 and Cu(I), and yield binding energetics comparable to experimental dissociation constants. The observed changes in the copper coordination environment might be the triggering signal for the transmission of the ethylene response.

## Introduction

The gaseous hormone ethylene regulates the growth and development of plants and acts in agronomically relevant processes, such as fruit senescence, ripening, and decay. In *Arabidopsis thaliana*, the main response to the hormone is mediated by a family of receptors composed of ETR1, ETR2, ERS1, ERS2, and EIN4 [1], which, in their functional state, form dimers and higher-molecular weight oligomers at the ER membrane [2, 3]. The overall structure of the receptor family has a modular arrangement that resembles bacterial two-component receptor histidine kinases, with an N-terminal transmembrane sensor domain, a GAF domain in the middle portion, and a catalytic transmitter domain at the C-terminus [4, 5]. It has been shown that the N-terminal transmembrane domain requires copper as a cofactor to functionally sense ethylene and trigger the response of the protein [6]. C65 and H69 have been found to be essential to protein function and are suggested to be part of the copper binding site [6, 7]. While the structure of the cytoplasmic part of the ETR1 receptor has been characterized experimentally and also by homology modeling [8–10], only recently we described the first *ab initio* model of the transmembrane domain of the protein, showing that two copper ions in the +1 oxidation state are bound with high affinity to the receptor [11]. This structural model is supported by a recent site-directed spin labeling and electron paramagnetic resonance spectroscopy study [12]. Although residues C65/H69 and copper are required for ethylene binding, and it is generally assumed that ethylene interacts with the copper ion, these interactions have yet to be directly observed. The ETR1 environment of one cysteine and one histidine per copper ion does not conform to the active site binding motifs typically found in copper proteins [13]. However, we note that from a molecular perspective that copper has rather flexible interaction possibilities, with coordination numbers that range from two to five [14]. These features of copper and the non-canonical environment could be essential to accommodate the binding of ethylene and regulate receptor activation. We thus set out to characterize the Cu(I) active site in the absence and presence of ethylene with XAS and EXAFS spectroscopy and by molecular dynamics simulations at a mixed quantum mechanics/molecular mechanics (QM/MM) level. The findings are consistent with the coordination of residues C65 and H69 to the Cu(I) site and exhibit changes upon addition of ethylene that are attributed to modifications in the Cu(I) coordination environment.

## Materials and Methods

### EXAFS and XANES experiments

The transmembrane domain region of ETR1 (ETR1^1-157^) was expressed and purified from *E. coli.*as previously described. The apo protein was loaded with monovalent copper using BCA2-Cu(I) [11]. Samples of ETR1 and ETR1 treated with ethylene (ETR1+ethylene) were loaded into custom Delrin X-ray sample cells with a 38 micron thick Kapton tape window and frozen and stored in liquid nitrogen until measurement. The total protein concentration of the measured sample was approximately 0.4 mM; [Cu] ~ 0.4 mM, in buffer S (50 mM TRIS/HCl pH 8 with 200 mM NaCl and 0.015% w/v Fos-Choline16) and additional 20% v/v glycerol as a glassing agent.

Partial fluorescent yield (PFY) Cu K-edge XAS data were recorded at the SSRL beamline 9-3 using a 100-element solid-state Ge detector (Canberra) with a SPEAR storage ring operating at ~500 mA and 3.0 GeV, as previously described [15]. The incoming X-rays were selected using a Si(220) double-crystal monochromator, and a Rh-coated mirror was utilized for harmonic rejection. Samples were maintained at ~10 K in a liquid helium flow cryostat. Data were calibrated by simultaneously measuring a copper foil, with the first inflection point set to 8980.3 eV.

Individual PFY-EXAFS scans were evaluated and processed in Matlab 2017a to average selected channels of the multi-element detector and normalize the averaged PFY signal to the incident beam intensity (I0). Final averaged scans were then further processed within Athena [16], where a second-order polynomial was fit to the pre-edge region and subtracted from the entire EXAFS spectrum. A three-region cubic spline (with the AUTOBK function within Athena) was employed to model the background function to *k*= 12 Å^-1^. Fourier-transforms were performed over a windowed *k*-range of 2 to 11.0 Å^-1^ and are presented without a phase shift correction.

Theoretical EXAFS spectra were calculated using Artemis utilizing the multiple scattering FEFF6 code.[16] The EXAFS amplitude, χ(*k*), is given by

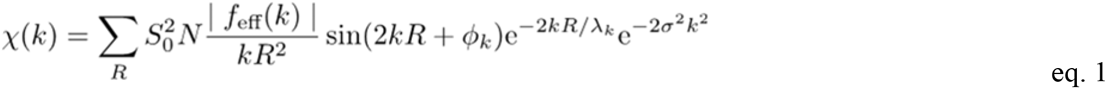

where 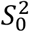 is the overall many-body amplitude factor, *N*, is the degeneracy of the path(s), |*f*_eff_(*k*)| is the effective scattering amplitude, and *R* is the absorber-scatterer distance. A Debye-Waller-like factor, exp(−2σ^2^*k*^2^), is also included to account for disorder. Lastly, *λ_k_* is the mean free path of the photoelectron, and *ϕ_k_* is the total photoelectron wave phase shift for the interaction between the absorber and the scatterer.

Individual scattering paths were calculated and fit within Artemis from FEFF6.[16] The Fourier-transform spectrum of each sample was fit over a range of *R* = 1.0 to 3.5 Å (non-phase shift corrected). The FT is the product of a transform of *k*^3^-weighted EXAFS spectrum with a Hann window over the range of *k* = 2 to 11.0 Å^-1^. By grouping similar scattering paths of a common coordination shell and increasing its degeneracy, *N*, the number of variables used for that coordination shell is minimal, 2 variables: σ^2^ and Δ*R*. A single *E_0_* variable was used for all paths in a given fit. *S*_0_^2^ was set to 0.9 for all paths. The goodness of final fits was evaluated by their reduced *χ^2^* as calculated within Artemis.

### Molecular dynamics simulations of ETR1 TMD, Cu(I), and ethylene systems

To generate a model describing changes in the ETR1 copper binding site upon ethylene binding, QM/MM MD simulations and a minimal structural model of the copper binding site were used. Recently, we performed MD simulations at the classical mechanical level of the model of the ETR1 TMD together with one Cu(I) per monomer and free diffusing ethylene to identify potential ethylene binding sites [11]. Here, we performed MD simulations of a model of the ETR1 TMD with ethylene and Cu(I) using a 12-6-4 Lennard-Jones ion model, which describes ion-solvent interactions more accurately [17], considering C65 and H69 as relevant for copper binding [6]. H69 was protonated at the epsilon nitrogen (HIE) to favor interactions of the delta nitrogen (ND1) with copper; C65 forms favorable interactions with the ion in a deprotonated state (CYM) [13]. These protonation states are consistent with a neutral pH environment and the *pKa* of Cu(I)-binding cysteines [18, 19]. From those simulations, all structures with a Cu-SG(Cys65) and Cu-ND1(His69) distance < 2.5 Å and a distance < 8 Å between an ethylene molecule and the copper ion were selected and clustered according to the ethylene orientation with respect to C65 and H69. The centroid of the most populated cluster was used as starting point for setting up QM/MM MD simulations, where residues C65 and H69 including their backbone atoms, Cu(I), and ethylene are included within the QM layer.

From the structure at the centroid, residues 62 to 72 of helix 2 of the TMD model were extracted to describe a minimal Cu(I)/ethylene binding site. The helix was embedded in a truncated octahedral TIP3P water box [20] with a distance of 14 Å to the box boundaries and parametrized using the force field ff14SB to describe the protein residues [21] and a 12-6-4 Lennard-Jones ion model to describe the copper in the molecular mechanics part [17], resulting in an overall charge-neutral system.

The assembly was minimized in five repeated mixed steepest descent/conjugate gradient minimizations with a maximum of 20,000 steps each, using positional restraints with a force constant of 25 kcal mol^-1^Å^-2^ on the protein residues, copper, and ethylene in the first step, reducing it to 5 kcal mol^-1^Å^-2^ in the intermediate steps, and removing all restraints in the last step. The relaxation was started by thermalizing the system from 10 to 100 K for 5 ps under NVT conditions, and from 100 to 300 K for 115 ps under NPT conditions at 1 bar, using an isotropic Berendsen barostat with a relaxation time of 1 ps. In all steps, the temperature was maintained by using Langevin dynamics, with a friction coefficient of 1 ps^−1^.

After completing 5 ns under these conditions, an initial QM/MM MD simulation was performed, where residues C65 and H69 including their backbone atoms, Cu(I), and ethylene were represented using the PM6 implementation in AMBER20 [22]. A half-harmonic restraint to a distance < 6 Å with a force constant of 0.1 kcal mol^-1^Å^-2^ between copper and the center of mass of ethylene was used to maintain ethylene in the proximity of Cu(I). After running the MD simulations for 1 ns, ethylene was pulled towards the Cu(I) using an adaptive steered approach [23], a force constant of 100 kcal mol^-1^Å^-2^, and a constant pulling speed of 1 Å ns^-1^ in 25 rounds with five independent replicas of 100 ps. The distance between the centers of masses (COM) of ethylene and Cu(I) was used as the reaction coordinate, starting from a distance of 4.14 Å. To obtain an energetically favorable transition path, after each round, the trajectory with the work profile closest to the Jarzynski average was selected [23], until the reaction coordinate reached a value of 1.74 Å.

### QM/MM US simulations of ETR1 helix 2, Cu(I), and ethylene

From the intermediate steps of the selected trajectories, 25 equidistant structures covering the reaction coordinate from 1.74 to 4.14 Å were used to set up umbrella sampling (US) simulations at a mixed QM/MM level. For this, the initial reaction coordinate values were restrained by harmonic potentials with a force constant of 100 kcal mol^-1^Å^-2^. To obtain significant sampling in a reasonable amount of time, the GPU implementation of TeraChem and a B3LYP 6-31G level of theory were used [24–26]. Again, residues C65 and H69 including their backbone atoms, Cu(I), and ethylene were considered in the QM part. Adaptive QM/MM (adQM/MM) simulations were applied, with a set of four replica exchange simulations per window and considering the six closest water molecules to the Cu(I) as the adaptive layer. To ensure the convergence of the adQM/MM calculations, the hybrid DIIS/A-DIISb scheme [27] with the “watcheindiis” flag in TeraChem was used [28]. Each window was sampled for 20 ps under adQM/MM conditions, which due to the replica exchange effectively represents 80 ps per window, recording structures every 100 fs and the reaction coordinate every 20 fs. The obtained distributions have a median overlap between windows of 42.6% (see below). The distributions were analyzed with the weighted histogram analysis method (WHAM) implementation from Alan Grossfield [29] using 100 bins between 1.8 and 4.2 Å, a tolerance of 10^-9^, and applying a Jacobian correction of 2RTln(*r*), with *r* being the reaction coordinate value [30]. The error of the free energy profile was computed as the standard error of the mean of independent measurements on 1 ps fractions of the data.

### Calculation of the ethylene binding equilibrium constant

To calculate the equilibrium constant from the free energy profile, a single dimension approximation was used, integrating along the reaction coordinate according to eq. 2 [31]:

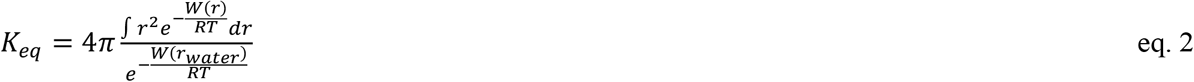

where *W* is the PMF value, *r* is the reaction coordinate, *r_water_* is the reaction coordinate with both binding partners separated in bulk water, *R* is the gas constant, and *T* is the absolute temperature. *W*(*r_water_*) was defined as the average of the WHAM bins at the ten farthest reaction coordinate values. To compare with experimental values, a standard state of 1 M was used by considering a factor of *C*°=1/(1661 Å^3^). From here, the standard free energy can be calculated as (eq. 3):

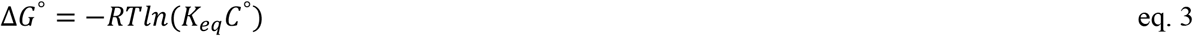

### Geometrical measurements from US distributions

Geometrical changes between bound and unbound states along the US simulations were evaluated. For this, the unbound state was defined as all structures lying at a reaction coordinate value higher than the PMF local minimum at 3.95 Å (see below). For the bound state, the energy minimum found at 1.89 Å (see below) was used as a reference, selecting all structures at ± 0.05 Å of the minimum, corresponding to the distance between US windows. Windows that showed the unbinding of the cysteine in the bound states were filtered out, considering only those where the SG-Cu(I) distance was below 3.89 Å, identified as the upper limit of the main peak of the distance distribution. All geometrical measurements were performed with CPPTRAJ [32].

### QM/MM minimization and vacuum optimization using B3LYP DEF2-TZVP with ORCA

To obtain a minimized/optimized structure, a representative structure of the bound and unbound states was obtained by clustering the filtered frames, using as a discriminating measure the root mean square deviation (RMSD) of the residues of the helix 2 peptide included in the QM region, i.e., C65, H69, ethylene, and Cu(I). The clustering was performed with CPPTRAJ [32], using the DBSCAN algorithm, a minimum of 4 points, and an epsilon parameter of 0.7 Å and 0.5 Å for the unbound and bound frames, respectively. The main cluster representative was either minimized using the AMBER/ORCA interface[33, 34] with B3LYP DEF2-TZVP[35] (SI Table 2) or optimized in vacuum with ORCA at the same level of theory (SI Table 3). For the latter, the input XYZ structure was obtained as generated by the AMBER interface by running a dummy 0 length calculation, and by constraining the N and C atoms of the termini of the considered C65 and H69 residues, to avoid major position changes as observed in the helix structure. Additionally, and to avoid sampling the bound state of ethylene in the unbound structure, a constraint between Cu(I) and each ethylene C atom was considered.

## Results

### Copper XAS

The Cu K-edge X-ray absorption spectrum of ETR1 (**Figure 1**) is typical of a Cu(I) site. No weak 1→3d pre-edge feature is observed that would be indicative of Cu(II). The first intense feature at 8983.5 eV is a Cu 1 s→4p transition of a Cu(I) ion, and the absorption energy of this feature is consistent with either a non-linear two or a three-coordinate copper center.[36] The Cu XAS of ETR1 in the presence of ethylene (ETR1+ethylene) exhibits an increase in the feature at 8983.5 eV. Previous studies of small molecules have established that normalized Cu(I) intensity of the 1s→4p feature of three-coordinate complexes is generally less than two-coordinate centers (approximately ~0.6 vs. 1.0 a.u., respectively). However, caution must be taken as the intensity of this 1 s→4p feature is modulated in three-coordinate complexes as the geometry is varied from T-shaped to trigonal planar [36]. Ultimately, the increase of this feature at 8983.5 eV for ETR1 + ethylene suggests a modification in the copper coordination environment in the presence of ethylene. In addition, the white line of the ETR1 samples at approximately 8995 eV exhibits a slight intensity decrease upon the addition of ethylene, providing further evidence for modulation of the copper environment in the presence of ethylene.

**Figure 1.**
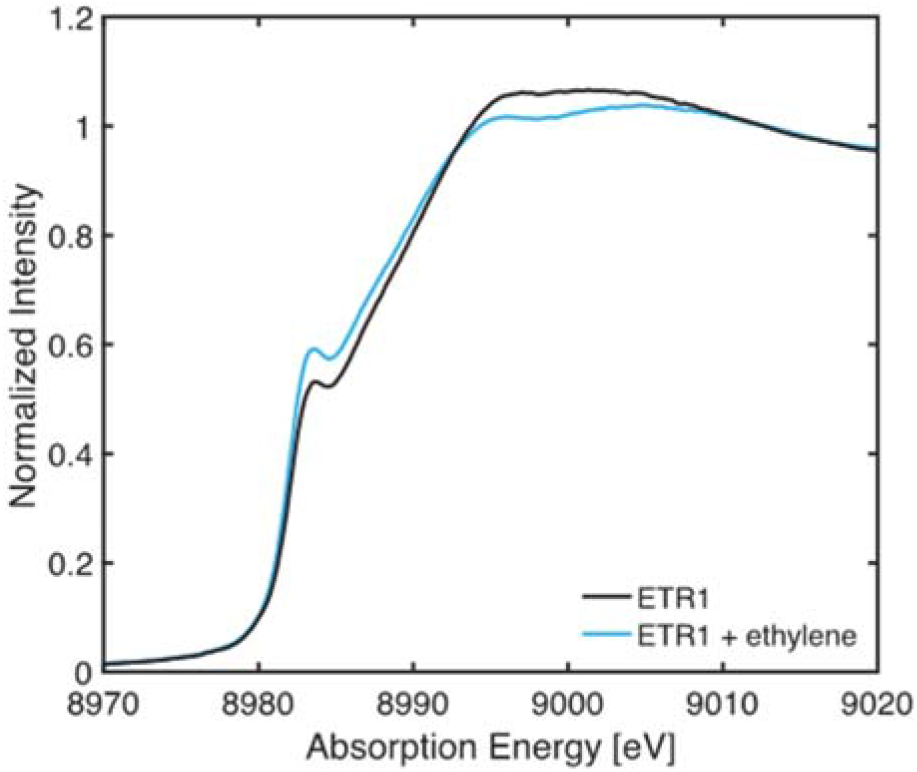
Normalized Cu K-edge XAS of ETR1 and ETR1 + ethylene.

### EXAFS

The *k*^3^-weighted EXAFS spectrum of ETR1 exhibits a single dominant waveform in the range of 2 to 8 Å^-1^ with a maximum amplitude at ~ 5 Å^-1^, **Figure 2a**. A similar EXAFS pattern is observed for ETR1 + ethylene, with a slight increase in intensity of the EXAFS signal at higher *k*. The non-phase shifted Fourier Transform of the EXAFS spectra over a *k*-range from 2 to 11 Å^-1^ yields a single asymmetric feature for each sample centered at a radial distance of *R* ~ 1.7 Å for ETR1 and slight shift to a longer radial distance of ~1.75 Å for ETR1 + ethylene, **Figure 2b**. While a phase-shift may be applied to the FT spectrum to approximately estimate Cu-X bond distances, fitting of the EXAFS below yields more precise Cu-X distances, as well as ligand identity information.

**Figure 2.**
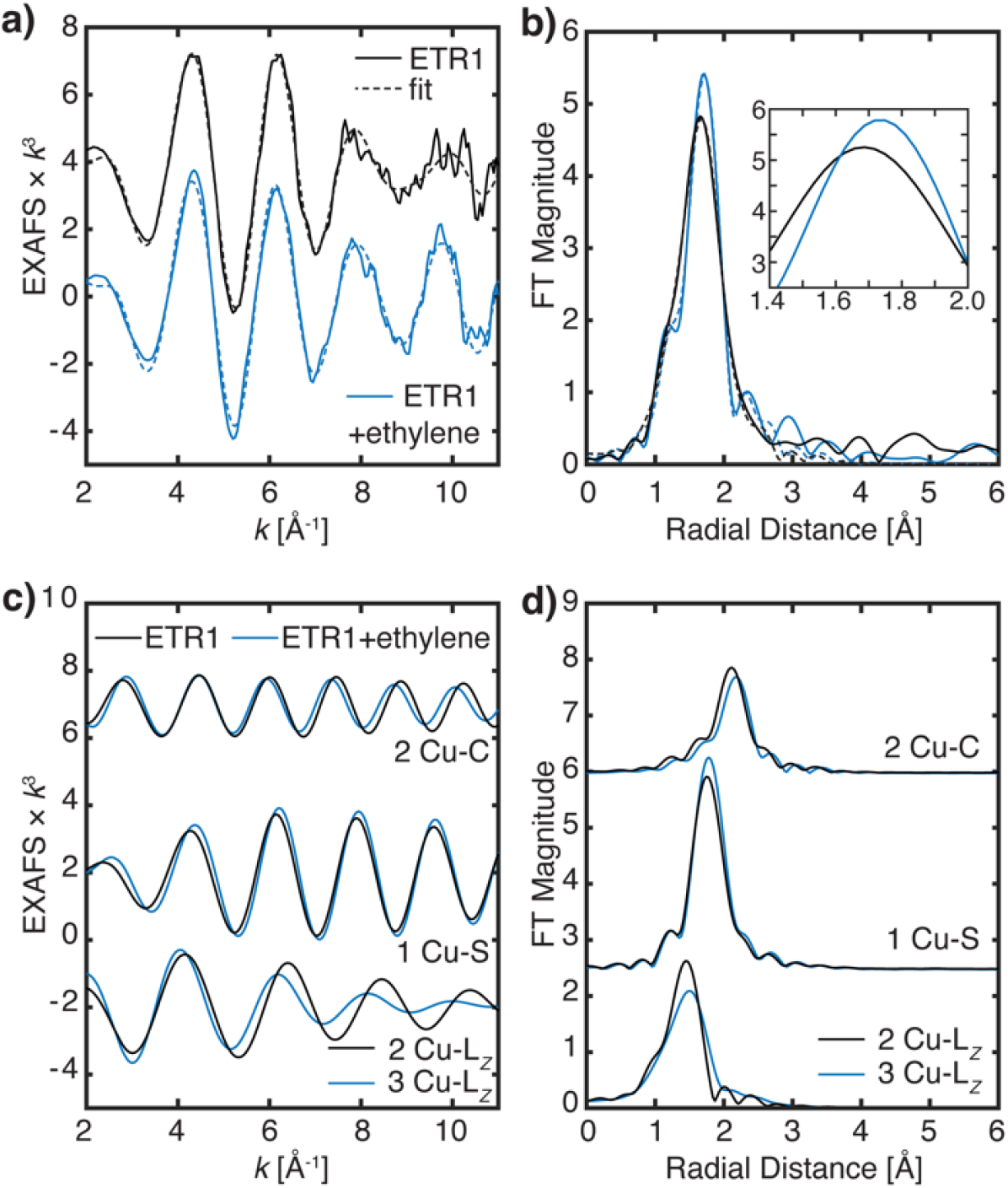
(a) *k*^3^-weighted EXAFS and (b) non-phase-shifted Fourier transform spectra of ETR1 and ETR1 + ethylene (solid lines) are plotted with each of their best fits, Fit 2 and Fit 6, respectively (dashed lines). (c, d) Individual scattering paths of fitted EXAFS for ETR1 (solid lines) and ETR1 + ethylene (dashed lines); the individual path groups have been vertically offset for clarity. The degeneracy of the Cu-C*long* and Cu-S paths are the same for each sample, however, for the Cu-L_*Z*_ path, ETR1 is fitted with *N* = 2 and ETR1 + ethylene with *N* = 3. The EXAFS spectra in panel a have been smoothed with a boxcar filter, *n* = 3 points.

The EXAFS of ETR1 is well fit with mixed light atom scatters (*L_Z_* = C, N, or O) and S in the first coordination shell, **Figure 2**. In a minimal fit with these two interactions (**Fit 1, Table 1**) at distances of 2.02 Å for Cu-*L_Z_* and 2.31 Å for Cu-S, these two scattering paths interfere constructively around k ~ 5 Å^-1^ yielding the maximal EXAFS amplitude, and are out-of-phase with respect to one another (deconstructive interference) at higher *k*, resulting in a loss of EXAFS amplitude, **Figure 2c**. The Fourier transform of *k*^3^-weighted EXAFS shows a single broad feature centered at an approximate radial distance of *R* ~ 1.70 Å, **Figure S1**. The fit of the EXAFS data is significantly improved, as judged by the reduced χ^2^ statistic, with the inclusion of a doubly degenerate Cu-C*_long_* scattering interaction of approximately 2.6 Å (**Fit 2**). This may represent a non-coordinating amino-acid side chain or second-sphere interaction. **Fit 2** of ETR1 yields 2 L_*Z*_ and 1 S scattering interactions with slightly shorter distances compared to distances to **Fit 1**, due in part to the differences in their ΔE0 values. The Cu-Lz distances of 1.95 Å in **Fit 2** is more consistent with what has been previously observed in low-coordinate Cu(I) nitrogen coordination spheres.[37–39]. The σ^2^-value of the single Cu-S path (3.72 × 10^-3^Å^2^) in **Fit 2** is close to the ideal value of 3.00 × 10^-3^Å^2^ for a single scattering interaction [40]. For the case of Cu-L*z*, when a single scattering path within a fit model represents multiple potential scattering paths/ligands the static disorder (σ^2^) is increased. This may be due to the inequivalent mean scattering distances of the multiple scattering paths modeled in the single path, resulting in a larger σ^2^-value [41]. The σ^2^-value of the Cu-L_*Z*_ scattering path is large (7.71 × 10^-3^Å^2^), but within an acceptable range for the increased static disorder. Attempts to fit ETR1 with a two-coordinate ligand set, 1L_*Z*_1S (**Fit 3**), yields both a non-physical σ^2^-value, where the σ^2^ value is too small (≪3.0 × 10^-3^Å for a single scattering path) [40], and an increased χ^2^ value indicating a poorer goodness-of-fit. Furthermore, the increased χ^2^ value of **Fit 3** also indicates a statistically poorer fit. This failure to fit the EXAFS with a two-coordinate environment further suggests a three-coordinate environment, as also suggested from the XAS edge analysis above. A four-coordinate fit, 3L_*z*_1S (**Fit 4**) has only a moderate improvement compared to **Fit 2**, indicated by the slightly decreased χ^2^ value. However, the XAS edge suggests a lower coordination number than four, leading us to favor **Fit 2**.

**Table 1.** Fitted EXAFS parameters of ETR1

| Fit | N | Path | R (Å) | ± | $\sigma^2$ ($10^{-3}$ Å<sup>2</sup>) | ± ($10^{-3}$ Å<sup>2</sup>) | $\Delta E_0$ (eV)<sup>a)</sup> | $\chi^2$ |
| --- | --- | --- | --- | --- | --- | --- | --- | --- |
| 1 | 2 | Cu-L<sub>Z</sub> | 2.017 | 0.020 | 4.203 | 1.458 | +6.80 | 253.74 |
|  | 1 | Cu-S | 2.305 | 0.021 | 5.679 | 1.932 |  |  |
| 2 | 2 | Cu- L<sub>Z</sub> | 1.948 | 0.017 | 7.705 | 1.326 | -2.73 | 58.63 |
|  | 1 | Cu-S | 2.220 | 0.014 | 3.722 | 0.786 |  |  |
|  | 2 | Cu-C<sub>long</sub> | 2.606 | 0.017 | 2.751 | 1.363 |  |  |
| 3 | 1 | Cu- L<sub>Z</sub> | 1.899 | 0.018 | 1.313 | 1.328 | -5.93 | 99.02 |
|  | 1 | Cu-S | 2.197 | 0.013 | 1.381 | 0.761 |  |  |
|  | 2 | Cu-C<sub>long</sub> | 2.587 | 0.018 | 0.775 | 1.217 |  |  |
| 4 | 3 | Cu-L<sub>Z</sub> | 1.991 | 0.020 | 11.955 | 1.674 | +0.24 | 49.58 |
|  | 1 | Cu-S | 2.233 | 0.018 | 5.972 | 1.329 |  |  |
|  | 2 | Cu-C<sub>long</sub> | 2.621 | 0.018 | 4.183 | 1.806 |  |  |

a) $E_0 = 8987$ eV

For the favored **Fit 2**, the long-range Cu-C/N scattering interactions of the imidazole group and other multiple-scattering paths are not clearly observed in the EXAFS and Fourier transform spectra. We note, however, that the relatively large σ^2^-value associated with the Cu-L_*Z*_ scattering path, coupled with the modest FT window available, suggests that well-ordered contributions from the imidazole ring may not be anticipated. Lastly, attempts to fit these longer scattering paths did not yield large statistical improvements in the fits and introduced a significant number of additional parameters.

The EXAFS model of ETR1 resolves a Cu-S scattering interaction at 2.22 Å that is consistent with Cu-S(Cys) coordination[37]. This distance is shorter than typical Cu-S(Met) distances[42]. Previously, ethylene binding studies of mutant ETR1 variants suggested that H69 and C65 are important residues for ethylene activity and form the potential site of copper binding [6, 7]. The mixed coordination sphere revealed by EXAFS agrees with the proposed coordinating residues based on previous structure-function studies and molecular mechanics studies [1, 7]. While the EXAFS model yields a total coordination number of three, potential water binding or an additional amino acid may not be conclusively assigned from this data. The analysis of the EXAFS and Cu K-edge together are consistent with a low-coordinate Cu(I) site.

Upon the addition of ethylene, the copper’s coordination environment is clearly modified as indicated by the changes in the Cu EXAFS presented in **Figures 2 and S2**. The intensity of the first coordination radial shell in the Fourier Transform spectrum (**Figure 2b**) increases and the peak’s maximum shifts to a longer radial scattering distance by +0.05 Å. The increase in intensity, as well as the shift to a longer radial distance, are both indicative of an increase in the copper coordination number – as the coordination number increases, the average metal-ligand bond lengths also increase. To understand the factors contributing to the observed changes in the EXAFS of ETR1 in the presence of ethylene, the data were first modeled using the same coordination sphere as ETR1 and then varied to fully elucidate the changes induced by the addition of ethylene (**Table 2**).

**Table 2.** Fitted EXAFS parameters of ETR1 +ethylene.

| Fit | *N* | Path | *R* (Å) | ± | $\sigma^2$ (Å<sup>2</sup>) | ± (Å<sup>2</sup>) | $\Delta E_0$ (eV)<sup>a)</sup> | $\chi^2$ |
| --- | --- | --- | --- | --- | --- | --- | --- | --- |
| 5 | 2 | Cu-L<sub>Z</sub> | 1.984 | 0.031 | 11.986 | 2.951 | -1.15 | 450.28 |
|  | 1 | Cu-S | 2.219 | 0.014 | 2.568 | 0.901 |  |  |
|  | 2 | Cu-C<sub>long</sub> | 2.645 | 0.026 | 3.654 | 1.807 |  |  |
| 6 | 3 | Cu-L<sub>Z</sub> | 2.025 | 0.031 | 16.967 | 3.410 | +0.95 | 397.94 |
|  | 1 | Cu-S | 2.223 | 0.012 | 3.200 | 1.025 |  |  |
|  | 2 | Cu-C<sub>long</sub> | 2.659 | 0.026 | 4.024 | 1.775 |  |  |
| 7 | 1 | Cu-L<sub>Z</sub> | 1.933 | 0.031 | 4.242 | 2.653 | -3.55 | 603.71 |
|  | 1 | Cu-S | 2.211 | 0.015 | 1.166 | 0.887 |  |  |
|  | 2 | Cu-C<sub>long</sub> | 2.627 | 0.027 | 2.711 | 1.901 |  |  |
| 8 | 3 | Cu-L<sub>Z</sub> | 2.086 | 0.031 | 8.028 | 4.106 | +8.95 | 1580.48 |
|  | 1 | Cu-S | 2.259 | 0.029 | 9.844 | 6.880 |  |  |

a) $E_0 = 8987$ eV

Fitting of ETR1 + ethylene with a 2L_*Z*_1S, 2C-long model (**Fit 5**) yields a comparable fit to to the favoured fit of ETR1 (**Fit 2**). The Cu-S scattering distances are equivalent in both samples, however, a smaller σ^2^-value for the ETR1 + ethylene **Fit 5** is found. The relatively decreased σ^2^-value of ~2.6 × 10^-3^ Å^2^ can represent a potentially more ordered Cu-S scattering interaction than in the ETR1 sample. The Cu-L_*Z*_ interaction fits to a slightly longer distance of 1.98 ± 0.03 Å, but is within error of the fitted distance of 1.95 ± 0.02 Å for ETR1 without ethylene. However, the σ^2^-value of the *N* = 2 Cu-L_*Z*_ path for ETR1 + ethylene fit, 12.0 (± 3.0) x 10^-3^ Å^2^, is significantly larger than the same path for ETR1, 7.7 (± 1.0) x 10^-3^ Å^2^. This indicates that the static disorder of the Cu-L_*Z*_ is increasing in the ETR1 + ethylene sample, despite the observation of the more ‘well-ordered’ Cu-S path. **Fit 5** does not explicitly consider any new Cu-C scattering interactions introduced by the addition of ethylene. The addition of ethylene to the copper’s first coordination sphere would introduce additional Cu-C scattering interactions at possible scattering distances in the range of 1.95 to 2.16 Å [43–47]. However, the utilized *k*-range of 2-11 Å^-1^ in the EXAFS (**Figure 2a**) yields a resolution of Δ*R* ~ 0.17 Å in the Fourier Transform spectrum (**Figure 2b**). This means that the Cu-C contribution from an interacting ethylene cannot be separately resolved from the Cu-L_*Z*_ scattering path, requiring the scattering interactions of the model to be further grouped. This should increase the fitted coordination number and, depending on the differences of the mean scattering distances of the individual paths in the group, further increase the static disorder of the Cu-L_*z*_ path.

**Fit 6** of ETR1 + ethylene with a 3-L_*Z*_1S, 2C-long (analogous to **Fit 4** for ETR1 without ethylene, see **Table 1**), where the first coordination shell number is increased, yields an improved fit by the evaluation of the χ^2^ value. Here, the σ^2^-value of the Cu-S scattering path is closer to the previously fitted σ^2^-value of ETR1 (**Fit 2**), implying no significant changes in the Cu-S disorder. Furthermore, the fitted Cu-S distances of the two samples are in excellent agreement with each other, indicating no significant differences. Within **Fit 6**, the *N* = 3 Cu-L_*Z*_ scattering path has a further increased σ^2^-value, which reflects the increased static disorder as expected for the additional Cu-C(ethylene) interaction(s). While this σ^2^ value may appear large, the Cu-L_*Z*_ modeled path nevertheless makes a real observable contribution to the fit as seen in **Figure 2c,d** and the σ^2^-value is still in an acceptable tolerance and for the increased coordination number, *N*. It is important to note that the coordination number, *N*, and σ^2^-values are negatively correlated, making it difficult to definitively assign a coordination number by EXAFS (i.e., 2 vs. 3). Inspection of the Fourier transform spectra reveal that the general intensity of the fitted Cu-S shell does not change between the two samples, however, a slight decrease for the Cu-L_*Z*_ shell does occur. This decrease of intensity is attributed to the increased disorder and broadening of the radial shell. A modest shift of the Cu-L_*Z*_ scattering distance is also observed, meaning the fitted mean Cu-L_*Z*_ scattering distance is longer in **Fit 6** for the ETR1 + ethylene sample, supporting the presence of a longer Cu-C(ethylene) interaction that increases the mean scattering distance of the Cu-L_*Z*_ path. Lastly, attempts to fit the EXAFS of the ETR1 + ethylene sample to a two-coordinate model (**Fit 7**) results in statistically poor fit, and the inclusion of the longer-range *Cu-C_long_* is also neccessary to improve the overall fit (**Fit 8**), similar to what was observed to ETR1.

Holistic comparisons between the fitted parameters of ETR1 (**Fit 2**) and ETR1 + ethylene (**Fit 6**) exhibit both an increased mean scattering Cu-L_*z*_ distance and increased static disorder for the scattering path upon the addition of ethylene to the sample, supporting the interaction and coordination of ethylene to the copper center of ETR1. The overall trend of marginally increased bond lengths, increased disorder, and the observed geometric distortion from the XAS edge upon the addition of ethylene to the sample are all consistent with the coordination of ethylene to the copper site of ETR1.

### Free energy profile of ethylene binding to and structural changes of the ETR1-Cu site

To obtain a structural model of the ETR1-Cu binding site to compare to the EXAFS data upon ethylene binding, adQM/MM US simulations of the binding process were performed. An ETR1 helix comprising residues 62 to 72, including C65 and H69 as main Cu binding partners, was embedded in explicit solvent. This simplified structural model only includes part of the overall protein and no membrane environment, leaving residues less constrained than under experimental conditions. US in 25 windows was performed, with the distance between the COMs of ethylene and Cu(I) used as the reaction coordinate, ranging from 1.74 to 4.14 Å. The computed potential of mean force (PMF) for the overall process has a global minimum of −12.48 kcal mol^-1^ at a distance of 1.89 Å (Figure 3). The equilibrium constant *K*_eq_ (eq. 2) of 4.7 × 10^6^ M^-1^ was computed by integration along the reaction coordinate, and from it Δ*G*°_binding_ = −9.11 ± 1.04 kcal mol^-1^ (eq. 3). Previously, dissociation constants for ETR1-Cu-ethylene in the range of (*K*_d_ = 0.04 - 1.24 μL L^-1^) were reported [48, 49]. Considering an ethylene density of 1.16 g L^-1^ at 20°C [50] and a molecular weight of 28.1 g mol^-1^ for ethylene, the experimental equilibrium constants relate to a *K*_eq_ = 2.0 × 10^7^ M^-1^-6.1 × 10^9^ M^-1^, and Δ*G*°_binding_ = −9.95 - −12.00 kcal mol^-1^. Hence, the computed *K*_eq_ is within chemical accuracy of 1 kcal mol^-1^ of the higher bound of the experimental values [51], lending support to the accuracy of the PMF computations.

**Figure 3.**
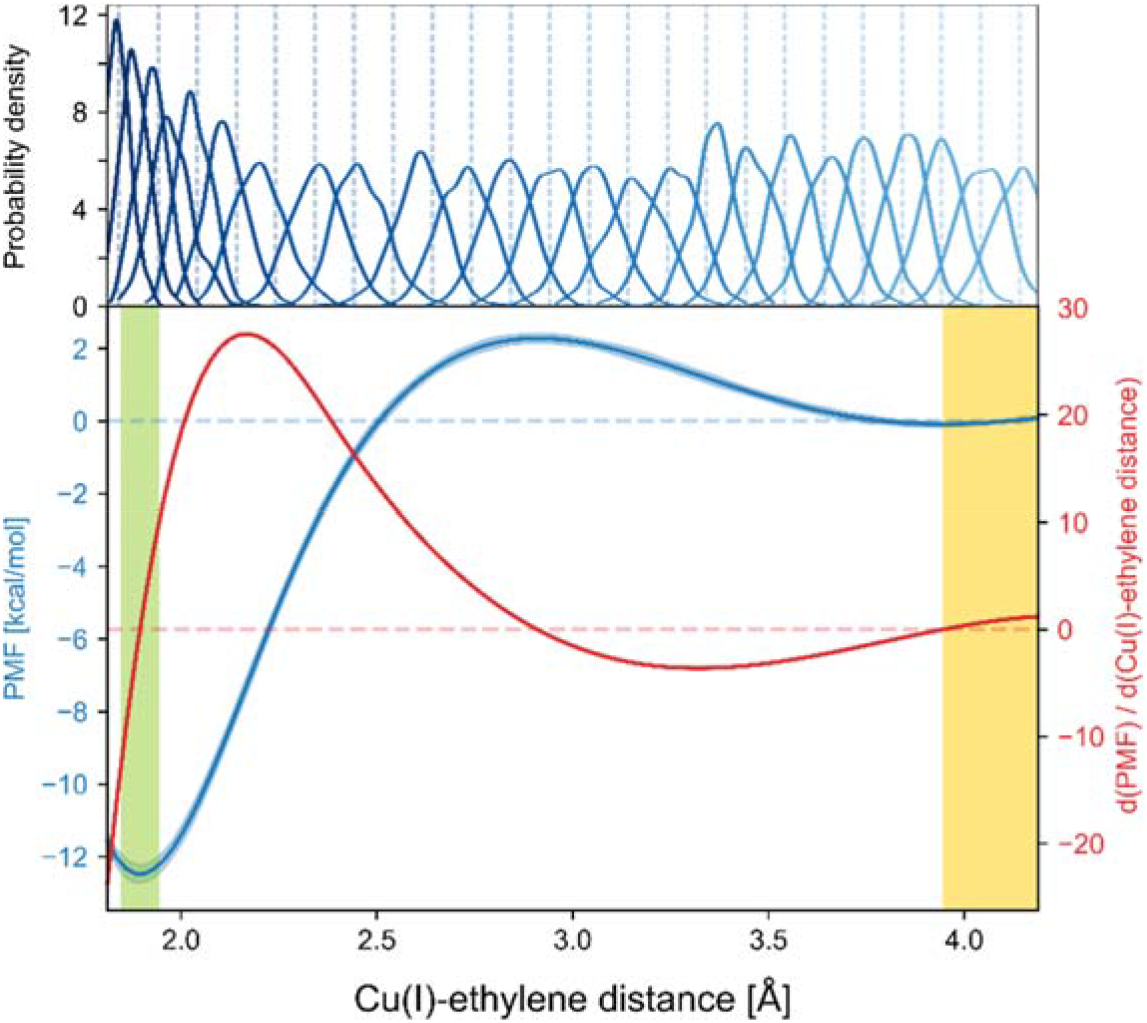
US and potential of mean force (PMF) of ethylene binding to Cu(I) in the ETR1 binding site, using the helix model. (Top) The distance between the center of mass of ethylene with respect to Cu(I) was used as a reaction coordinate, sampling 25 windows with a median overlap to adjacent windows of 42.6%. (Bottom) Potential of mean force (PMF) obtained from WHAM setting the value for unbound ethylene to zero (blue), and its first derivative (red). A global minimum is observed at 1.89 Å (12.48 kcal mol^-1^). The free energy barrier has a maximum at 2.90 Å. The reaction coordinate values at the minima were used to categorize bound (green area) and unbound (yellow area) states. The blue shaded area shows the standard error of the mean.

To probe for the binding site geometry in the unbound and bound states, structures where C65 and H69 form interactions with the Cu(I) ion were selected from the US simulations (see Materials and Methods, Figure 3 yellow and green shaded areas for unbound and bound states). Consistent with the Dewar-Chatt-Duncanson model [52], the hybridization states of the ethylene carbon atoms change upon interacting with the Cu(I) ion, leading to an elongation of the C=C bond from 1.34 Å to 1.40 Å (experimentally, the CC-distance in etyhlene is 1.313 Å [53] and in ethane 1.409 Å [54]) and an improper dihedral angle involving HHC=C of 159° rather than 180°, revealing an out-of-plane movement of the hydrogen atoms (Figure 4a and 4b, SI Table 1). Crystallographic and spectroscopic analyses revealed a C=C distance between 1.33 and 1.37 Å in ethylene-Cu complexes [55–58]; distances between 1.37 and 1.42 Å were previously reported in DFT studies [59–62]. At the global minimum of the PMF, the distance between Cu(I) and the ethylene carbons is 2.00 Å (SI Table 1), as previously reported [59–62].

**Figure 4.**
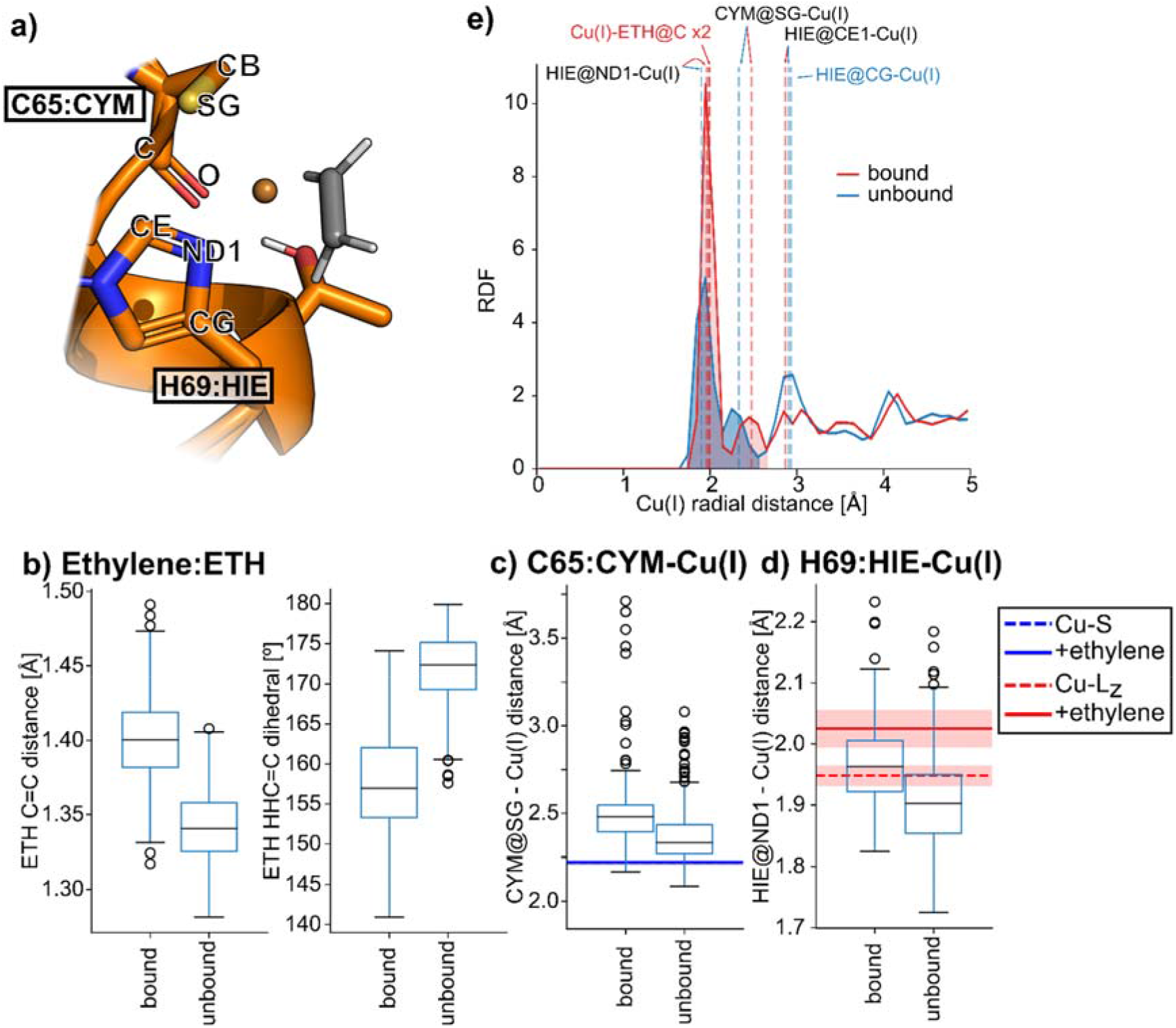
Geometric properties of the Cu(I) binding site and ethylene in unbound and bound states of ETR1 (helix model as shown in a) obtained from US simulations. b) The C-C distance of ethylene increases upon binding, and the coordination at the carbons becomes less planar, indicating a change in the hybridization state, in agreement with the Dewar-Chatt-Duncanson binding model [63]. Upon ethylene binding, the distances between Cu(I) and C65 (CYM) S_γ_ (CYM@SG) (c) and H69 (HIE) N_δ_ (HIE@ND1) (d) increase. All median distances can be found in SI Table 1. The blue/red lines and shaded areas indicate the expected distances and standard deviations of Cu-S and Cu-L_Z_ as obtained from the best EXAFS fits with and without ethylene (Tables 1 and 2). For distances to the closest carbon atoms on both residues, see Figure S3. e) Heavy-atom radial distribution function for Cu(I) calculated from the bound and unbound states. The shaded area corresponds to a coordination number of ~3 and ~4 for the ethylene unbound and bound states, respectively. Dashed lines show atoms with a median distance < 3 Å, as shown in panels c), d), Figure S3 and SI Table 1.

Upon ethylene binding, the distance between Cu(I) and S_γ_ of C65 increase by 0.15 Å (Figure 4c, SI Table 1). Three out of 25 windows along the reaction coordinate showed the unbinding of C65; these frames were not considered in the analysis. The EXAFS measurements do not exhibit such a large increase in the distance to the sulfur atom upon ethylene binding (the distances of Cu-C*_long_* and Cu-L_Z_ increase by 0.03-0.07 Å (Tables 1 and 2); see also below). No Cu-C distance computed here to carbons in the helix distinctly match the Cu-C*_long_* distance from EXAFS models (Figure S3). Note, however, that some of the carbons in the binding site environment of the full protein might not be included in our helix model and/or the Cu-C_*long*_ path in the EXAFS model may represent multiple indiscriminate scattering interactions that are not resolved at the resolution of the experiment. The experimental change in the distance between Cu and N_δ_ of H69 is not uniquely resolvable from other light atom contributions (including the coordinated ethylene), as discussed above. Nevertheless, the general trends in the computed Cu-N(His) distance, in the presence and absence of ethylene, may be considered consistent with the EXAFS trends (Figure 4d).

The general trend of increasing distances between S_γ_ of C65 and N_δ_ of H69 to Cu(I) upon ethylene binding is also obtained when using a larger basis set in a B3LYP DEF2-TZVP QM/MM minimization employing the AMBER/ORCA interface and in an ORCA optimization in vacuum only considering residues C65, H69, and Cu(I), starting from cluster representatives of the filtered frames (see Methods, and SI Tables 2 and 3). Interestingly, while both calculations show a lower distance between S_γ_ of C65 and Cu(I) in the unbound state (closer to the distance obtained from the EXAFS fit, see Table 1), there is a marked increase in this distance when ethylene is bound to Cu(I).

The Cu(I) coordination number as computed from the radial distribution function is 4.22 for the bound states, corresponding to N_δ_ of H69, ethylene, and S_γ_ of C65 (Figure 4e and SI Table 1), while for the unbound states, a coordination number of 2.95 was computed, corresponding to N_δ_ of H69, S_γ_ of C65, and a water molecule. The computed increase of the first coordination shell number agrees with the increase of the *L_Z_* coordination number assigned in the EXAFS fitting upon the addition of ethylene and the suggested coordination environment change from the XAS. In addition, the increase in the distance observed for *L_Z_* upon ethylene binding agrees with the formation of the interaction between Cu(I) and ETH@C (see Tables 1 and 2 and SI Table 1).

To conclude, the binding free energy computed from the dissociation constant obtained from the free energy profile computed with aQM/MM US simulations agrees within chemical accuracy (< 1 kcal mol^-1^ [51]) with experimental data, supporting the chosen molecular model and the simulation setup. Computed changes in the ethylene geometry upon binding correspond with hybridization changes expected from the Dewar-Chatt-Duncanson binding model. Our results indicate that N_δ_ of H69 relates to the Cu-L_*Z*_ scattering path of fitted EXAFS. The computed distance change to S_γ_ of C65 agrees qualitatively with measured data to a sulfur atom, although the computed magnitude is too large; this difference might originate from insufficient constraints within our helix model or differences in the experimental (10 K) versus computational (300 K) temperatures used. The change in the coordination number of Cu(I) upon ethylene binding from C65, H69, and a water molecule to C65, H69 and ethylene might be a trigger for signal transmission within the ETR1 TMD that can lead to receptor inhibition.

## Discussion

We have shown that the Cu(I) ion of ETR1 is a low-coordinate copper center with a mixed first coordination sphere of light L_*Z*_ and heavier S ligands. The fitted Cu-S EXAFS distance assigns a Cu-cysteine interaction, likely with C65, while the shorter Cu-L_*Z*_ interaction could be explained by a histidine ligand, likely H69. By QM/MM US simulations of ethylene binding to a minimal ETR1 Cu-binding site, we obtained a *K_eq_* of 4.7 × 10^6^ M^-1^ with Δ*G*°_binding_ = −9.11 ± 1.04 kcal mol^-1^, in very good agreement with values measured experimentally. In the simulations, the ethylene C=C bond distance increases, and the HHC=C improper dihedral angle decreases, upon binding to Cu(I), akin to hybridization changes predicted by the Dewar-Chatt-Duncanson model. Upon binding of ethylene, the coordination of Cu(I) changes by increasing the interaction lengths with H69 and C65, and replacing a water molecule with ethylene.

Distances between Cu(I) and coordinating atoms computed from the simulations for the ethylene-bound and -unbound states overall increase, in qualitative, but not quantitative, agreement with the EXAFS measurements. According to the distance, the Cu-L_*Z*_ interaction assigned from EXAFS is consistent with H69 being one of the interacting ligands in the ETR1 binding site. The imidazole ring contributes to the coordination of Cu in the bound and unbound states, whereas the thiolate moiety of C65, suggested to be strongly coordinating to Cu(I) in the absence of ethylene[64], is shifted towards longer distances upon ethylene binding (Figure 4e), in agreement with observations that suggest a preference for an imidazole ring in single-point QM DFT calculations [65]. This coordination changes might be the triggering signal for the transmission of the ethylene signal in the sensor domain, which then is propagated to the C-terminal region of ETR1. Alternatively, in the same study [65], Cu(I) was suggested to be additionally coordinated by one of the Tyr residues present in the binding site, probably Y32 located on helix 1 of ETR1; helix 1 was not included in our simulations. We cannot rule out the possibility of this contributing to the Cu-L_Z_ scattering path of the EXAFS, as the suggested Cu-O distance to the hydroxyl oxygen of Y32 of 1.89 Å (see model M2 in [65]) falls within the resolution of the separable scattering interactions of the Fourier transform of the EXAFS.

In summary, C65 and H69, previously described as critical residues for ethylene binding [6, 7], likely coordinate to the Cu(I) in ETR1. Furthermore, our combined XAS, EXAFS, and QM/MM US studies demonstrate that ethylene interacts with the transmembrane domain of ETR1 and modulates the coordination sphere of the Cu(I) center. How exactly the binding of ethylene causes the inactivation of ETR1 remains to be determined, but we hypothesize that either the change in coordination of Cu (I) or steric influences of the bound ethylene, or both, lead to a signal transmission within the TMD towards the ETR1 C-terminus located in the cytoplasm.

## Acknowledgements

This study was funded by the Deutsche Forschungsgemeinschaft (DFG, German Research Foundation) project no. 267205415 / CRC 1208 grant to GG (TP B06) and HG (TP 03). GEC and SD thank the Max Planck Society for financial support. Use of the Stanford Synchrotron Radiation Lightsource, SLAC National Accelerator Laboratory, is supported by the U.S. Department of Energy, Office of Science, Office of Basic Energy Sciences under Contract No. DE-AC02-76SF00515. The SSRL Structural Molecular Biology Program is supported by the DOE Office of Biological and Environmental Research, and by the National Institutes of Health, National Institute of General Medical Sciences (P30GM133894). The contents of this publication are solely the responsibility of the authors and do not necessarily represent the official views of NIGMS or NIH. We are grateful for computational support and infrastructure provided by the “Zentrum für Informations- und Medientechnologie” (ZIM) at the Heinrich Heine University Düsseldorf and the computing time provided by the John von Neumann Institute for Computing (NIC) to HG on the supercomputer JUWELS at Jülich Supercomputing Centre (JSC) (user ID: HKF7, VSK33, ETR1).

