## Supporting Information for "Spectroscopic and QM/MM studies of the Cu(I) binding site of the plant ethylene receptor ETR1"

#### Supplementary Figures

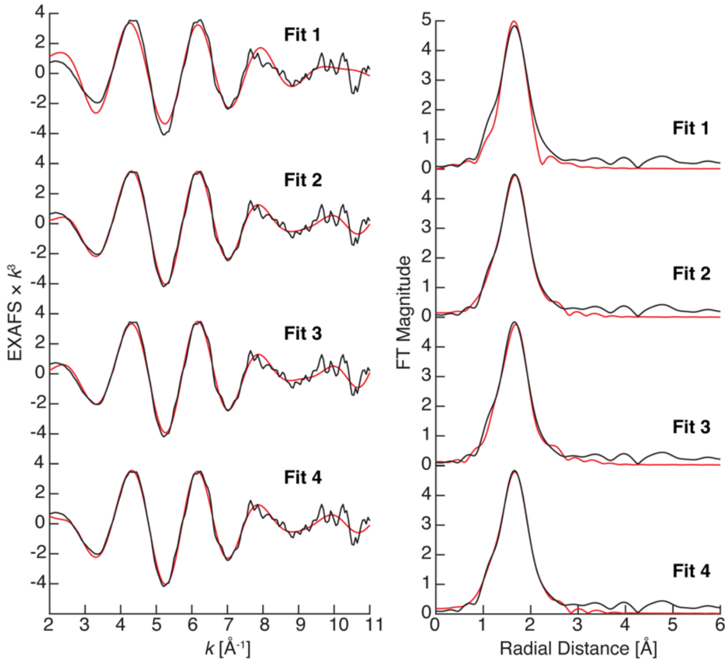

**Figure S1.** (left) $k^3$-weighted EXAFS and (right) non-phase-shifted Fourier transform spectra of ETR1 (black) and fitted spectra (red). The raw EXAFS spectra (left) have been smoothed with a boxcar filter, $n = 3$ points. Fit parameters are detailed in **Table 1**.

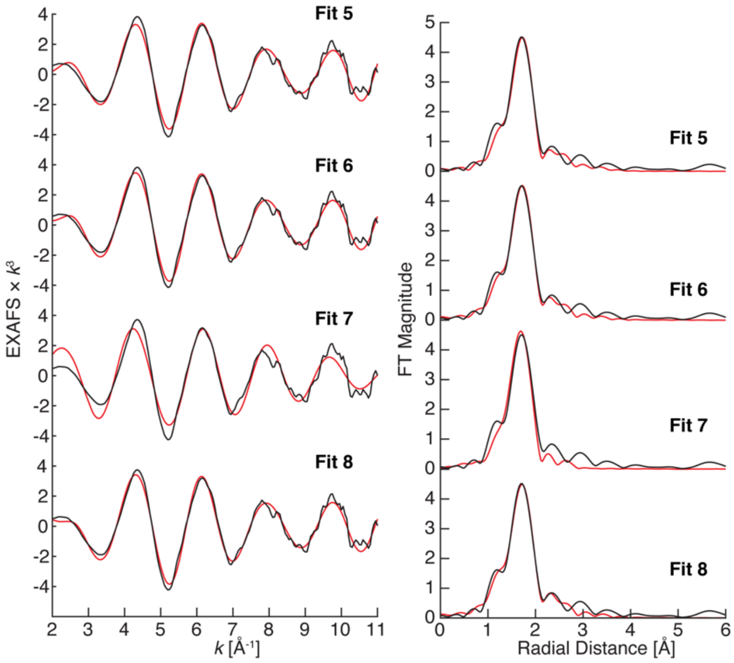

**Figure S2.** (left) $k^3$-weighted EXAFS and (right) non-phase-shifted Fourier transform spectra of ETR1 + ethylene (black) and fitted spectra (red). The raw EXAFS spectra (left) have been smoothed with a boxcar filter, $n = 3$ points. Fit parameters are detailed in **Table 2**.

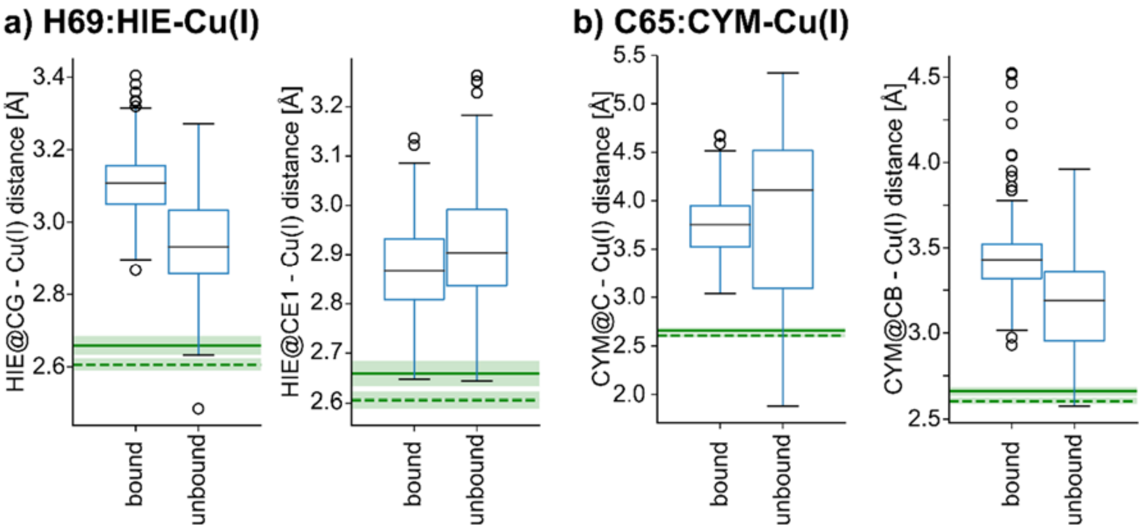

**Figure S3.** Distances of Cu(I) in the binding site with unbound and bound ethylene to closest C atoms of H69 and C65 obtained from US simulations. a) Upon ethylene binding, the distances between Cu(I) and H69 (HIE) Cγ (HIE@CG) increase, whereas the distance to Cε (HIE@CE1) decreases. b) For C65 (CYM), the distances to the carbonyl carbon (CYM@C) fluctuate less in the bound state than in the unbound state, while the distance to Cβ (CYM@CB) increases. Median distances can be found in SI Table 1. The green lines and shaded areas indicate the expected distances and standard deviation of Cu-$C_{long}$ as obtained from the best EXAFS fits with and without ethylene (Tables 1 and 2).

### Supplementary Tables

**SI Table 1.** Median geometric properties ± median absolute deviation of the median obtained from unbound and bound states of ETR1 (helix model) and Cu(I) from QM/MM US simulations at the B3LYP 6-31G level using TeraChem[1]

| **Property** | **unbound (*N* = 479)** | | **bound (*N* = 299)** | |
|---|---|---|---|---|
| | | M.A.D.[c] | | M.A.D.[c] |
| ETH C=C distance [a] | 1.34 | 0.02 | 1.40 | 0.02 |
| ETH HHC=C dihedral [b] | 172.4 | 2.97 | 158.3 | 4.20 |
| Cu(I)-ETH@C distance [a] | 4.35 | 0.35 | 2.00 | 0.03 |
| Cu(I)-ETH@C1 distance [a] | 3.93 | 0.35 | 1.99 | 0.03 |
| CYM@C-Cu(I) distance [a] | 4.11 | 0.48 | 3.76 | 0.20 |
| CYM@O-Cu(I) distance [a] | 3.92 | 0.61 | 3.26 | 0.31 |
| CYM@CB-Cu(I) distance [a] | 3.19 | 0.20 | 3.43 | 0.11 |
| CYM@SG-Cu(I) distance [a] | 2.33 | 0.08 | 2.48 | 0.08 |
| HIE@CG-Cu(I) distance [a] | 2.93 | 0.09 | 3.11 | 0.06 |
| HIE@ND1-Cu(I) distance [a] | 1.90 | 0.05 | 1.96 | 0.04 |
| HIE@CE1-Cu(I)distance [a] | 2.90 | 0.08 | 2.87 | 0.06 |

[a] In Å.

[b] In °.

[c] Median absolute deviation

**SI Table 2.** Optimized geometric properties obtained from bound and unbound states of ETR1 (helix model) and Cu(I) from QM/MM cluster representatives at the B3LYP DEF2-TZVP level using the AMBER/ORCA interface [2, 3]

| **Property** | **unbound** | **bound** |
|---|---|---|
| ETH C=C distance [a] | 1.33 | 1.36 |
| ETH HHC=C dihedral [b] | 178.9 | 167.7 |
| Cu(I)-ETH@C distance [a] | 5.03 | 2.13 |
| Cu(I)-ETH@C1 distance [a] | 4.17 | 2.14 |
| CYM@C-Cu(I) distance [a] | 3.74 | 3.87 |
| CYM@O-Cu(I) distance [a] | 3.47 | 3.66 |
| CYM@CB-Cu(I) distance [a] | 3.05 | 3.38 |
| CYM@SG-Cu(I) distance [a] | 2.19 | 2.45 |
| HIE@CG-Cu(I) distance [a] | 2.99 | 3.14 |
| HIE@ND1-Cu(I) distance [a] | 1.92 | 2.01 |
| HIE@CE1-Cu(I)distance [a] | 2.88 | 2.85 |

[a] In Å.

[b] In °.

**SI Table 3.** Optimized geometric properties obtained from ethylene bound and unbound states of residues C65, H69, and Cu(I) from cluster representatives at the B3LYP DEF2-TZVP level using ORCA[2], constraining the N- and C-termini of the included residues.

| **Property** | **unbound** | **bound** |
|---|---|---|
| ETH C=C distance [a] | 1.33 | 1.37 |
| ETH HHC=C dihedral [b] | 179.5 | 167.3 |
| Cu(I)-ETH@C distance [a] | 4.73[c] | 2.13 |
| Cu(I)-ETH@C1 distance [a] | 3.72[c] | 2.07 |
| CYM@C-Cu(I) distance [a] | 3.58 | 3.43 |
| CYM@O-Cu(I) distance [a] | 3.67 | 3.48 |
| CYM@CB-Cu(I) distance [a] | 3.19 | 3.29 |
| CYM@SG-Cu(I) distance [a] | 2.16 | 2.30 |
| HIE@CG-Cu(I) distance [a] | 3.04 | 3.22 |
| HIE@ND1-Cu(I) distance [a] | 1.94 | 2.06 |
| HIE@CE1-Cu(I)distance [a] | 2.87 | 2.89 |

[a] In Å.

[b] In °.

[c] Distance constrained during optimization
